# ioSearch: a tool for searching disease-associated interacting omics; application on breast cancer data

**DOI:** 10.1101/2022.08.01.502395

**Authors:** Sarmistha Das, Deo Kumar Srivastava

## Abstract

Biomarkers identification is difficult for cancer and other polygenic traits because such complicated diseases occur due to an intricate interplay of various genetic materials. Although high-throughput data from recent technologies provide access to a tremendous amount of information still there is a huge gap in harnessing knowledge from the generated multi-omics data. It is evident from the availability of subject-specific multi-omics data from large consortiums that there is a growing need for appropriate tools to analyze such data. Traditional single-omics association tests more often identify strong signals but fail to explore the between-omics relationship and find moderately weak signals due to multiple testing burdens. Multi-omics data integration intuitively provides a clear advantage in understanding the genetic architecture of disease a little better by imparting complementary information. But the construction of such methods is challenging because of the diversity in the nature of multiple omics and the sample size which is much less than the number of omics variables. It is important to consider factors such as data diversity and prior biological knowledge to make meaningful and better predictions. Dimension reduction techniques such as feature selection are used to circumvent the sample size issue in general but treating all the omics variables similarly might be an oversimplification of the complex biological interactions. The lack of appropriate approaches for biomarker identification from complex multi-omics data led us to develop this method. ioSearch is a tool for integrating two omics assays with continuous measurements. Based on a two-step model, ioSearch explores the inter-relationship of the omics in a principal regression framework and selects features using sparse principal component analysis to provide easily interpretable inference in terms of p-values. Also, it uses prior biological information to reduce multiple testing burdens. Extensive simulation results show that our method is statistically powerful with a controlled type I error rate. Application of ioSearch to two publicly available breast cancer datasets identified relevant genes and proteins in important pathways.

## Introduction

Identifying genes and proteins responsible for the aberrant proliferation rate of cancer cells is essential for developing gene-based cancer therapy or drug development ^1,2^. For example, in breast cancer, state-of-the-art multi-gene panel screenings for germline/somatic pathogenic variants in the coding regions are more often advocated to determine the treatment regime ^3^ for the patients. Some of the genes in the panel are putative driver genes harbouring driver mutations. But the efficacy of a screening panel for detecting breast cancer signals is still unclear ^4^. Since any cancer is highly heterogeneous, it is difficult to detect its signal based on some fixed mutation set. This leads to the availability of clinical management guidelines only for patients with definitively actionable mutations ^5^. One reason may be that only some hereditary cases occur due to mutations in rare but highly penetrant genes, while most other genes in the testing panels are moderate-penetrance genes with low prevalence ^6^. A number of low-penetrance risk variants have been identified over years but the effect sizes of such individual single-nucleotide polymorphisms (SNPs) are quite small ^6^. Besides, given the fact that only 2% mutations are related to the protein-coding genome and non-coding regions may harbour driver genes ^7^, to detect or elucidate cancer prognosis we clearly require more downstream information.

It is now known that cancer cells require a sustained activation of protein expression to initiate and proliferate.This leads to abnormal protein production that deregulates the activation of signaling pathways and produces proteins required for tumorigenesis in patients. These aberrant proteins may also facilitate cancer progression and/or aid the cancer cells to develop resistance to treatment^8^. Thus, translation deregulation that is, alteration in the levels of protein synthesized from its messenger RNAs (mRNAs) is a primary downstream output of oncogenic signaling. Besides, the translation of mRNAs into proteins is a key event in the regulation of gene expression ^9^. But various studies suggest that the correlation between mRNA and corresponding protein levels in a cell is more often weak ^10^. One reason is that it is still unclear how cancer cells prioritize the translation of certain mRNAs over others from a pool of competing mRNAs. While some studies indicate cancer cells prioritize the translation of certain mRNAs that have phenotypic hallmarks of malignancy ^9^, others suggest even in absence of actionable mutations at the DNA level, it is possible that cancer cells may prioritize the translation of certain mRNAs that encodes for components of a signaling pathway ^8^. Some animal models also highlight that pathway activity triggered by a mutation in a signaling pathway could confer higher oncogenic potency than a mutation in an oncogene itself ^11^. Furthermore, many pathways have been targeted for gene-based therapy development and experimentally silencing certain genes in these pathways has amplified the potential benefit of the treatment ^12^. Thus, pathway information may elucidate crucial insights into disease aetiology by functionally relating genes and gene products located far apart on the chromosome.

Identification of genes and gene products, on the other hand, is important for the development of gene-based therapies and also for finding causes of resistance to such therapies ^13^. But the experimental approach of gene silencing is expensive and requires a lot of knowledge base in that very focused area for conducting the experiments. So, it is experimentally infeasible to search for all aberrantly interacting genes and gene products (such as proteins) in the whole genome that either differentiate between disease status or various disease subtypes. Besides, even when the data generated from biological experiments is rich, smaller sample size may interfere with the statistical power of the experiments. In such scenarios, integrating information from larger consortium datasets and multi-omics data analysis may accumulate crucial prior information to minimize the list of target genes/gene products for further functional (eg. gene knockout or silencing) experiments. This might also reduce the growing gap between the ability to generate high-throughput omics data with the capacity to integrate, interpret, and harness the required knowledge to develop targeted drugs.

Nevertheless, the sample size is a permanent challenge in data integration because the number of omics features always exceeds. Yet numerous studies have focused on the integration of various omics data such as mRNA ^14^, DNA methylation ^15^, protein etc. to find disease-associated gene signatures. A lot of emphases has been put on integrating protein and mRNA expressions as they may provide complementary information on the disease ^16^. Recent technological developments such as RNA sequencing (RNA-Seq), reverse phase protein arrays (RPPA) etc. have paved the way for high-throughput data generation that provides unprecedented knowledge on a number of omics. Multi-omics data integration offers a clear advantage in accumulating information from these assays across different omics revealing crucial interactions unobserved by single-omics analyses. Hence, to integrate diverse omics data, some studies focused on experimental approaches ^17–19^ while others developed computational methods. Other than graph-based methods ^20^, among computational approaches, feature selection by canonical correlation or covariance ^21,22^, unsupervised learning ^23^ using factor analysis, supervised learning ^24^ using kernel functions based on prior knowledge (pathway information) have been studied. While most of the computational methods concentrate on dimension reduction through feature selection, the experimental methods ^17^ still focus on the most concordant pairs of genes and corresponding proteins. Some computational methods also apply spearman’s correlation to integrate transcriptome and proteome data and select the concordant pairs deregulated between normal and tumor individuals ^25^. This overlooks important information because, as already mentioned the relation between genes and their corresponding products (eg. proteins) is not straightforward (only weak to moderately correlated). The lack of a strong correlation has been conferred to a number of biological phenomena ^17,26,27^. On the other hand, feature selection methods considerably reduce the dimension of the data but the biological interpretation of a significant feature is complicated compared to the original variables.

In this article, we propose an integrated regression framework that selects features from heterogeneous omics data based on prior information from pathways and applies multivariate multiple regression to explore the inter-relation between the most deregulated variables between two subtypes or disease status in order to identify biomarkers. The advantages of this algorithm include prior-information-based dimension reduction of the entire omics data matrices (eg. transcriptome, proteome etc.), efficient handling of multicollinearity, and reduction of multiple testing burdens that minimize the chance of missing weaker signals compared to commonly used highly concordant gene-protein pair. The inclusion of pathway information also facilitates the exploration of long-range functional relations between and within different omics as opposed to single gene-based biomarker identification methods. Besides, a comprehensive testing algorithm for querying composed hypothesis ^28^ embedded in our algorithm generates conveniently interpretable p-values from an asymptotic distribution that reduces computation cost compared to permutation-based techniques. Simulation results show that our method is statistically powerful with a controlled type I error rate.

We applied our method to the protein and gene expression datasets of nearly 800 and 100 breast cancer samples respectively from The Cancer Genome Atlas (TCGA) and Clinical Proteomic Tumor Analysis Consortium (CPTAC) projects. Interestingly, we identified a list of genes and proteins most of which are reported to have interacting functional implications in breast cancer susceptibility across independent experimental studies. For example, in the AMPK signaling pathway, we identified PRKAA2 and AKT1S1(PRAS40) genes in both TCGA and CPTAC along with IRS1, IGF1R, CCND1, TSC1 etc. genes to be associated with disease status (early versus late-stage). But some of the important genes such as PRKAA2, IRS1 and IGF1R were unidentified by single-omics analyses. However, experimental studies have already confirmed (1) the role of PRKAA2 in the alteration of CCND1 expression as well as AMPK growth control and apoptotic signaling, (2) the correlation of PRAS40 phosphorylation with IGF1R-induced resistance to epidermal growth receptor inhibition in head and neck cancer, and (3) the effect of TSC1 in controlling insulin-PI3K signaling via regulation of IRS proteins. So, this method not only finds interacting multi-omics biomarkers but also captures weaker signals that are missed by traditional single-omics analyses. Thus, our algorithm ioSearch (interacting omics Search) holds a reasonable potential to be used as a tool to identify multi-omics biomarkers to narrow down the list of omics for further experimental analysis.

## Methods

### Overview of ioSearch

Suppose that we have two matrices *M* and *G* of the order *n*x*s*_*m*_ and *n*x*s*_*g*_ (Figure 1A) corresponding to protein and mRNA expression respectively where *n, s*_*m*_ and *s*_*g*_ denotes the sample size, number of genes and proteins. Based on pathway information (Figure 1B) (or any user-defined choices eg. functional annotations) ioSearch splits these matrices into smaller matrices (sub-matrices). A pair of such sub-matrices obtained one from each omic assay comprise a set (Figure 1C). Let us suppose in set *j* (*j* = 1, 2, …, *J*), we have 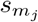 proteins and 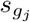 genes such that 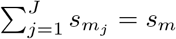 and 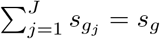. Next, we consider a two-step model to explore the joint effect of the interacting proteins and genes on the disease status for each set and finally combine the results from *J* sets to identify the disease-associated biomarkers.

**Figure 1:**
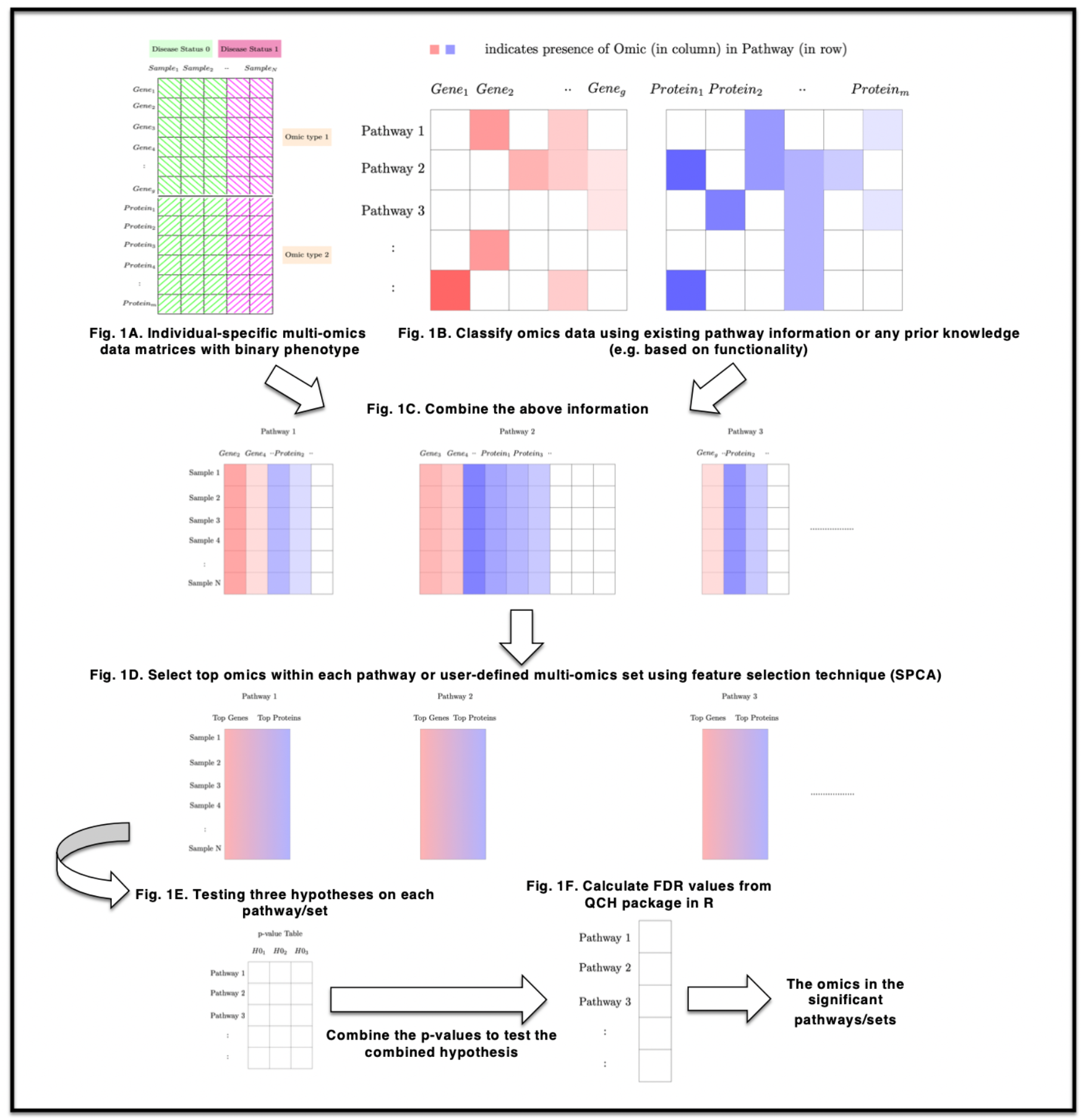
Graphical abstract.

In the first step, we model the effect of all proteins and genes in a set on the disease status using logistic regression. To do so, we regress the disease status on the principal components (PC) obtained from these genes and proteins. The reason behind employing PCs as opposed to original variables is to tackle any multicollinearity issue if it arises, avoid loss of degrees of freedom, and use maximum information contained in the variables. However, our demonstration is restricted to the first PC only, but it can easily be extended further. In mathematical notation, for individual *i*, the first step in our model can be written as,

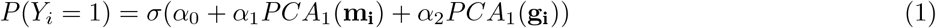

where, *Y*_*i*_, ***m***_*i*_, ***g****i*, and *PCA*_1_ denote the disease status, vector of 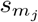 protein expressions, vector of 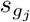 mRNA expressions, and first PC respectively. It is important to note that, even in a set/pathway, the number of genes and proteins could be large which may exceed the sample size. Moreover, it is much more likely that only some of these genes and proteins exhibit the strongest effects. So, the idea is to further explore the variables with maximum variance (based on sparse principal components loadings; See detailed algorithm Step 1) because they are expected to be most deregulated compared to the remaining variables in the set.

In the next step, the relation between the top (a user-defined number) proteins and genes that are selected based on top sparse PC loadings (Figure 1D) is further explored (See detailed algorithm Step 2). Let us consider the subsets 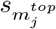 and 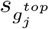 are respectively selected from 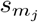 protein and 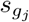 gene expressions based on the top sparse PC loadings. To establish whether any of these top omics are interacting among themselves, we perform a multivariate multiple regression of top proteins on top genes using a linear model as follows,

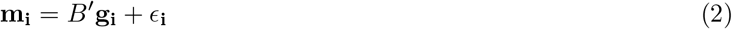

where 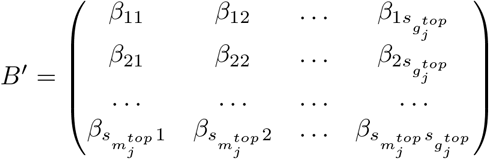 is the regression coefficient matrix of fixed components of order 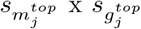. We assume 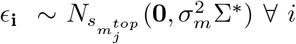 independently. Now, to jointly test if there is an association between the genes and proteins in the set, given that they play an important role in the association with the disease, we construct the hypothesis

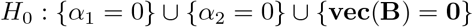

where 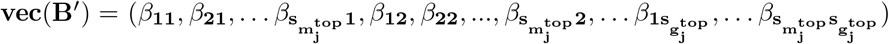 (See detailed algorithm Step 3). In the final step (See detailed algorithm Step 4), we use the R package ‘qch’ to test our composite hypothesis for all the sets together (Figure 1E-1F). So, the rejection of the null hypothesis corresponding to set *j* would mean the selected top omics in that set might jointly affect the disease status (i.e. at least one of the variables from each omics group interacts with the other and both the omic types are associated with the disease status). Thus, the resulting significant sets searched from among the entire transcriptome and proteome provide a collection of interacting proteins and genes that might differentiate between two subtypes or case-control status. Further, this testing method analyzes multi-omics profiles of the individuals together as opposed to combining results from two separate single-omics analyses.

### Detailed algorithm

Step 1: Although the interpretation of regressing all the omics on the disease status separately is easier, weaker signals will be lost due to multiple testing burdens. On the other hand, accumulating information from multiple omics will provide more information but directly using all the omics in the model will reduce the degrees of freedom. Moreover, if multicollinearity exists, then variances of regression coefficients may become very large. So, predictor variables are replaced by uncorrelated PCs. But retaining all the PCs is equivalent to using all the variables. We keep only the first PC because it contains the maximum information contained in each omics dataset (*M* and *G*) in any set/pathway.

Even when only one PC is retained, the interpretation is difficult because all the original variables remain in the PC. Keeping in mind, that only some omics might have larger effects we shrink some of the coefficients in the PC to zero using sparse PC analysis (SPCA) ^29^ so as to obtain the omics having the strongest effect sizes. SPCA uses the elastic net (EN) approach which provides a few advantages compared to ordinary lasso. First, is the high dimensional data handling. When the number of predictor variables exceeds the sample size (say, *N*), the ordinary lasso is restricted to a selection of at most *N* predictor variables, which is clearly unsatisfactory. But EN can potentially include all the variables in the fitted model. Moreover, EN handles the grouping effect, which means it tends to select a group of highly correlated variables once one variable among them is selected. But lasso restricts the selection to only one variable belonging to the same group. Since, in our context, it is more meaningful to include all the deregulated variables even though they are correlated or exceed the sample size, we employ SPCA to select a user-defined number of top omics.

Step 2: In the next step, we explore the between-omics relationship among the selected top omics to find interaction, if any. But again, multicollinearity might arise because the genes selected using SPCA although important, could be correlated. So, we use principal component regression (PCR) to estimate the regression coefficients. PCR framework allows ordinary least square estimation of the uncorrelated PC predictors guarding against large variance of the regression coefficients which is inevitable in presence of multicollinearity. To demonstrate the PCR framework, let us re-consider Eqn.(2) in matrix notation.

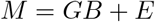

where, *E* = (*ϵ*_**1**_, *ϵ*_**2**_, … *ϵ*_**n**_)′ and *ϵ*_**i**_ ~ *N*_*t*_(0, Σ) where 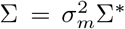 and 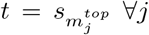. So, we may write the distribution of *M* as,

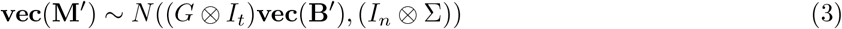

where **vec**(**M**′) is a vector that indicates all the elements of the matrix *M* have been stacked by rows. Thus, the likelihood function of this distribution can be written as,

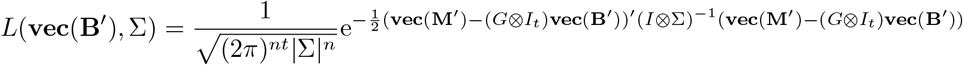

For simplicity let us assume, *G* ⊗ *I*_*t*_ = *G*^∗^ and **vec**(**B**′) = ***β***. Now let us define a matrix *Z* = (**z**_**1**_, **z**_**2**_, … **z**_**n**_) ′ to be the values of PCs for each individual. So, *Z* can be written as, *Z* = *G*^*∗*^*A*^*∗*^ where, 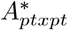 denotes the loading matrix whose *j*^*th*^ column is the *j*^*th*^ eigenvector of 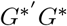 and 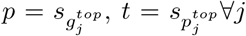. Because *A*^∗^ is orthogonal, *G* ^∗^ ***β*** can be written as, 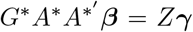 (say) where, 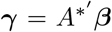. Finally, denoting **vec**(**M′**) as **M**^∗^ and **vec**(**E**′) as ***ϵ***^**∗**^ we have, ***M***^**∗**^ = *Z****γ*** + ***ϵ***^**∗**^. This simply replaces the original predictors by PCs in the regression model. In this work, we do not drop any PC for further dimension reduction. But appropriate choices could be made to select some PCs t|o furth|er reduce the dimension.

Step 3: To establish the connection between the previous two steps, we may consider the following approximate conditional distribution where the disease status of an individual *i* can be written as a function of the individual’s omics profiles,

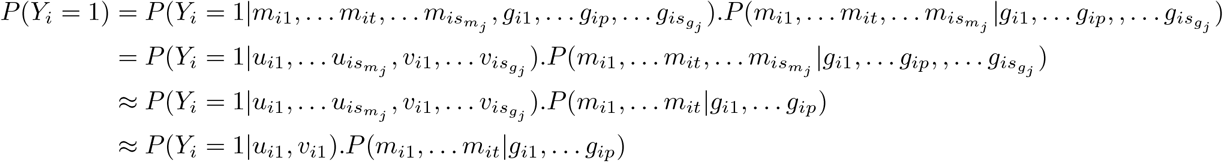

where, 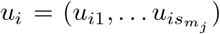 and 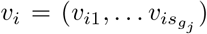 respectively denotes the PCs based on **m**_**i**_ and **g**_**i**_ corresponding to the *i*^*th*^ individual. The approximations are due to the fact that (1) SPCA only retains the omics with maximum variability (which are explored in Step 2) and (2) only the first PCs containing maximum information from each omics type are selected for the logistic regression (in Step 1). The rationale behind this step is that individual omic variables often may be too weak to provide any association signal with the disease. But together the top omics from different sources (based on loadings) are expected to provide more information. So, if the first PC of proteins and genes are associated with the disease status, it might some or all of the selected top omics are associated with disease because they are the most deregulated (having maximum variation) ones in that set/pathway.

Next, we write, the joint likelihood distribution of the above model as,

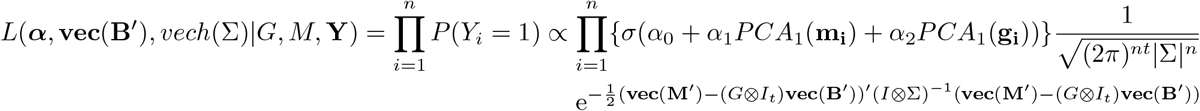

Then we estimate the parameters in the above function using maximum likelihood estimation. The estimates obtained by equating partial derivatives of the log-likelihood function (with respect to the parameters) to zero are as follows,

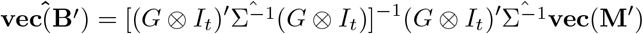

Since in Σ we assume only 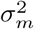 is the unknown parameter, while, Σ^*∗*^ is estimated from observed pairwise correlation of *t* proteins we estimate 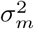using 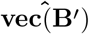 as,

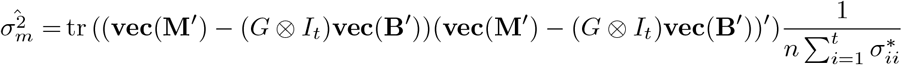

where, 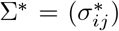. Moreover, since no closed form is possible for the estimates of ***α*** = (*α*_0_, *α*_1_, *α*_2_)′, we use the R package ‘glm’ to compute numerical values of the estimates.

Re-writing the PCR framework of Step 2 in matrix notation we get, *M* = *GB* + *E* = *GAA*′*B* + *E* = *W* Γ + *E* (say). Analogous to the estimate of **vec**(**B**′) we can write the estimate of **Vec(Γ′)** as, 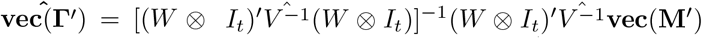 where, 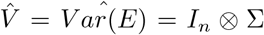. Now, using Γ = *A* ′*B* we can get 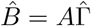 which implies, 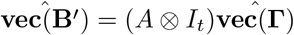

Step 4: Intersection-Union test (IUT) is a natural method for constructing a hypothesis test for our hypothesis of interest. According to IUT, the overall null hypothesis can be rejected only if each of the individual hypotheses can be rejected. Here, we use a testing method QCH ^28^ where IUT holds as a particular case. QCH is a scalable method that fits a mixture model to the p-values obtained from individual null hypotheses and the rejection rule is based on the posterior probabilities. Based under mild conditions, this method can perform efficiently for a large number of items such as pathways in our case. The advantage of employing the QCH testing method in ioSearch is that the entire set of omics data from multiple assays are combined in a single test to provide inference.

To test individual hypotheses, we construct test statistics for each of them. For testing *H*_01_ : *α*_1_ = 0 and *H*_02_ : *α*_2_ = 0, our test statistics will be 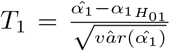 and 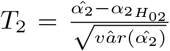 respectively. The test statistic for testing *H*_03_ : **vec(B′) = 0** will be 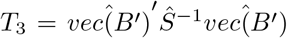 where, 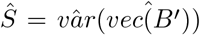 is the estimate of variance of 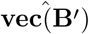. Under *H*_0_, *T*_1_ and *T*_2_ both follows *t*_*n*−1_ and *T*_3_ follows 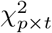.

## Results

### Simulation

We perform extensive simulations to study the performance (in terms of statistical power) of ioSearch for three different scenarios by generating two normally distributed omics say protein and gene expressions. Since these expression data are usually non-normal, ioSearch incorporates algorithms for the appropriate transformation of the datasets for real data analysis. However, simulating data involving randomly chosen real-life biological interactions might have an unknown confounding effect. So, we gradually increase the interaction complexity. For simplicity, in scenario 1, we assume interaction occurs between the expression of one gene and one protein. In scenario 2, we only increase the number of interacting genes to two such that, only one protein expression interacts with two gene expressions. Lastly, in scenario 3, we increase only the number of interacting proteins to two, implying that the expression of only one gene interacts with two protein expressions. To make the scenarios more realistic under each scenario, we assume a few more gene and protein expressions are associated with disease but do not interact with other omics. In case of the presence of more than one interaction as in scenarios 2 and 3, we demonstrate in detail the performance of ioSearch for different magnitude and directions of between-omics interactions.

To understand the effect of the number of sets based on prior information (eg. pathways) that are created from the protein and gene expression data matrices, we simulated all the scenarios for *J* sets each harbouring 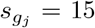 genes and 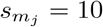 proteins where *J* is either 500 or 2000. Thus, it makes the total number of genes under study in the range of 7.5*K* to 30*K*, which is close to the number of data points obtained in RNA-Seq experiments. In our simulation, the total number of protein expressions varies from 5*K* to 20*K*. But, the number of identified known proteins is not that huge, but we still simulate such a dimension. This is to show that our algorithm is generic, meaning that we may replace proteomic or transcriptomic datasets with any other type of omics datasets having continuous measurement. Besides, the reason behind fixing the number of genes and proteins in each set is to avoid any extra weightage due to set size. This is not a criterion for the algorithm because ioSearch works when pathway sizes (in terms of the number of cognate gene and protein expressions) differ as shown in the applications section. But it might be better to split very large pathways into meaningful sub-pathways in real data analysis to capture more information. Ideally, only a few sets or pathways might harbour significant interacting gene and protein expressions in reality. So, we assume two situations where (1) 20% and (2) 10% sets to be significant under each simulation setup with sample sizes 100, 150, and 200. For every simulation, we assume an equal number of individuals in each of the phenotype classes to avoid confounding effects of any kind in the calculation of statistical power. But again, this is not a criterion for using the ioSearch algorithm as demonstrated in the real data application section where sample sizes in the disease classes are unequal.

For data generation, we simulate correlated protein expressions from a normal distribution with a mean as a function of the gene expressions if the protein interacts with the gene expression. The regression parameter between such interacting gene and protein expression is uniformly generated under different simulation setups. If any protein expression does not interact with the gene expression but is associated with disease status, the difference in mean between the disease status is assumed to be 1. In cases where protein expressions are neither disease-associated nor interacting with any gene expression, we assume means to be 0 for both the disease statuses. For the variance-covariance matrix, we assume a correlated structure with a common variance equal to 2. Gene expression data, on the other hand, are simulated independently from normal distributions with a fixed variance equal to 2 and a mean difference equal to 1 between disease statuses if the gene expression is assumed to be disease-associated, 0 otherwise. The variance is fixed for simplicity and to avoid any confounding effect from diverse variability between genes. Next, we generate phenotype data from a binomial distribution using Eqn 1. We assume, *α*_0_ = 1, *α*_1_ and *α*_2_ are uniformly generated under different simulation setups (indicated corresponding to each simulation result table). For the generation of omics data in the non-significant sets, we used different combinations of *α*_1_ and *α*_2_ and matrix *B* pertaining to the null hypothesis (i.e. at least one parameter is zero). For example, when 500 sets/pathways are simulated with 10% significant sets in the true model, we generated 50 significant sets and the remaining 450 sets from a variety of combinations viz. 90 sets (20% of the non-significant sets) with all zero parameter values (*α*_1_ = *α*_2_ = 0 and *B* = 0), 60 sets each (~ 13.5% of the non-significant sets) from the other combinations in the null hypothesis (i.e either one or two parameters with zero values). For generating significant sets, the mentioned parameters were assumed to be all non-zero with values satisfying different pre-defined scenarios. In case, when the percentage of significant sets is increased to 20% (i.e 100 significant sets among 500 sets), under the null hypothesis we generated 100 sets (25% of the non-significant sets) with all zero parameter values and the remaining 300 sets are generated equally from the rest (10%) of the combinations in the null hypothesis. We preserve the percentage of data generated in each category as cited in the above example when the number of sets/pathways is amplified to 2000.

From Tables 1 − 3 we observe that in each scenario the power of our method increases with sample size and does not depend on the number of sets/pathways. However, as expected the power increases as the % of true alternate sets increases, within a fixed number of sets/pathways. For scenarios 2 and 3, we simulated the between-omics interactions for a variety of magnitude and directions. In Table 1, the magnitudes differ greatly and the directions are opposite. In Table 2, the magnitudes are varied but not the directions. In Table 3, the directions are opposite keeping the magnitudes similar.

**Table 1:** Power calculation for ioSearch based on 100 replications. For all scenarios, *α*_1_ and *α*_2_ are selected from U(1,2) and U(0.3,0.5) respectively. For scenario 1 the regression coefficient of the interacting omics is generated from U(0.8,1). For scenarios 2 and 3, the two non-zero coefficients are selected from U(0.8,1) and U(−0.2,-0.1).

| No. of Pathways | % of true alternate sets | Scenario 1 |  |  | Scenario 2 |  |  | Scenario 3 |  |  |
| --- | --- | --- | --- | --- | --- | --- | --- | --- | --- | --- |
|  |  | Sample Size |  |  |  |  |  |  |  |  |
|  |  | 100 | 150 | 200 | 100 | 150 | 200 | 100 | 150 | 200 |
| 2000 | 20 | 0.4731 | 0.6825 | 0.8667 | 0.5551 | 0.8349 | 0.9420 | 0.4739 | 0.7029 | 0.8774 |
| 2000 | 10 | 0.4412 | 0.6275 | 0.8025 | 0.4850 | 0.73205 | 0.8980 | 0.4442 | 0.6304 | 0.7954 |
| 500 | 20 | 0.4586 | 0.7148 | 0.8532 | 0.5619 | 0.8328 | 0.9355 | 0.4576 | 0.7085 | 0.8528 |
| 500 | 10 | 0.4236 | 0.6464 | 0.7962 | 0.4662 | 0.7488 | 0.8926 | 0.4064 | 0.6388 | 0.7878 |

**Table 2:** Power calculation for ioSearch based on 100 replications. For all scenarios, *α*_1_ and *α*_2_ are selected from U(1,2) and U(0.3,0.5) respectively. For scenario 2, the regression coefficients of the interacting omics are generated from U(0.8,1) and U(0.1,0.2) while for scenario 3, they are simulated from U(0.8,1) and U(0.6,0.7).

| No. of Pathways | % of true alternate sets | Scenario 2 |  |  | Scenario 3 |  |  |
| --- | --- | --- | --- | --- | --- | --- | --- |
|  |  | Sample Size |  |  |  |  |  |
|  |  | 100 | 150 | 200 | 100 | 150 | 200 |
| 2000 | 20 | 0.5312 | 0.7260 | 0.8373 | 0.4132 | 0.7501 | 0.9157 |
| 2000 | 10 | 0.4135 | 0.7010 | 0.8257 | 0.4536 | 0.6685 | 0.8602 |
| 500 | 20 | 0.4995 | 0.7342 | 0.8526 | 0.4537 | 0.7559 | 0.9016 |
| 500 | 10 | 0.4402 | 0.7012 | 0.8244 | 0.4301 | 0.6554 | 0.8166 |

**Table 3:** Power calculation for ioSearch based on 100 replications. For all scenarios, *α*_1_ and *α*_2_ are selected from U(1,2) and U(0.3,0.5) respectively. For scenario 2, the regression coefficients of the interacting omics are generated from U(0.35,0.45) and U(−0.45,-0.35) while for scenario 3, they are simulated from U(0.8,1) and U(−0.7,-0.6).

| No. of Pathways | % of true alternate sets | Scenario 2 |  |  | Scenario 3 |  |  |
| --- | --- | --- | --- | --- | --- | --- | --- |
|  |  | Sample Size |  |  |  |  |  |
|  |  | 100 | 150 | 200 | 100 | 150 | 200 |
| 2000 | 20 | 0.4360 | 0.6192 | 0.7853 | 0.607 | 0.8009 | 0.9162 |
| 2000 | 10 | 0.4031 | 0.5903 | 0.7370 | 0.5763 | 0.7837 | 0.8911 |
| 500 | 20 | 0.4250 | 0.6347 | 0.7935 | 0.613 | 0.8301 | 0.9152 |
| 500 | 10 | 0.3776 | 0.5914 | 0.7474 | 0.5666 | 0.7876 | 0.8858 |

**Table 4.** Power of DIABLO based on 100 replications under three scenarios with same parameters as that in Table1

| No. of Pathways | % of true alternate sets | Scenario 1 |  |  | Scenario 2 |  |  | Scenario 3 |  |  |
| --- | --- | --- | --- | --- | --- | --- | --- | --- | --- | --- |
|  |  | Sample Size |  |  |  |  |  |  |  |  |
|  |  | 100 | 150 | 200 | 100 | 150 | 200 | 100 | 150 | 200 |
| 2000 | 20 | 0.4669 | 0.5025 | 0.5118 | 0.4814 | 0.5179 | 0.5128 | 0.4841 | 0.5208 | 0.5139 |
| 2000 | 10 | 0.4348 | 0.4859 | 0.5014 | 0.4544 | 0.5068 | 0.5036 | 0.4561 | 0.5099 | 0.5060 |
| 500 | 20 | 0.4673 | 0.5030 | 0.5139 | 0.4809 | 0.5070 | 0.5137 | 0.4851 | 0.5117 | 0.5163 |
| 500 | 10 | 0.4343 | 0.4857 | 0.5026 | 0.4532 | 0.4933 | 0.5050 | 0.4510 | 0.5007 | 0.5077 |

We did not find many data integration approaches other than DIABLO ^22^ that identify biomarkers to differentiate between disease statuses to compare with our method. DIABLO performs feature selection technique for dimension reduction in a supervised learning framework and is able to handle multiple omics. DIABLO focuses on biomarker identification and subtype classification on the basis of the selected features. In order to compare our method with DIABLO, we simulated datasets with parameters generated from the same distributions under each scenario. For DIABLO, as in our method, the power increases with sample size and also with the % of true alternate sets when the number of sets/pathways is fixed but it remains independent of the number of pathways. All three observations are in agreement with ioSearch. But the powers are much less than ioSearch and also the increase in power of DIABLO with sample size is very small.

### Applications

#### TCGA and CPTAC breast cancer overview

We applied ioSearch on 794 breast cancer (BC) patients from The Cancer Genome Atlas (TCGA) and 107 from Clinical Proteomic Tumor Analysis Consortium (CPTAC) patients. The TCGA dataset comprised female BC patients with ductal and lobular neoplasms belonging to the TCGA-BRCA project while the CPTAC dataset included female patients with invasive breast carcinoma. Although the type of invasive breast carcinoma is not mentioned in the CPTAC dataset, the most common types are invasive ductal carcinoma and invasive lobular carcinoma. Besides, ductal or lobular carcinoma in Stage 0 is generally considered non-invasive or pre-invasive. So, we excluded all Stage 0 patients from the TCGA dataset so that both the datasets may include similar disease types.

We applied our method to the transcriptome and proteome datasets obtained from both datasets. First, we removed the genes and proteins with missing data. Next, we applied Box-Cox and log transformations to normalize the protein and gene expressions. After removing outliers, we used Shapiro–Wilk test we check the validity of the normal assumption of our model. A few non-normal omics (≤ 5%) are removed prior to the data analysis. Thus, we obtained 4386 protein-coding genes and 222 proteins common in both datasets that intersected with KEGG pathway genes. Our phenotype of interest is the standard TNM (Tumor, Node, Metastasis) stage for breast cancer. The stage of each patient is based on the Tumor size, whether or not the tumor has spread into neighbouring (that is in the drainage area of the affected organ) lymph nodes, and if the tumor has spread to other parts of the body. We dichotomised the four stages into two by accumulating all types of Stages I (IA, IB) and II (IIA, IIB) into one subgroup and remaining in the other. This is because stages I and II are generally considered early-stage breast cancer while stage III is viewed as locally advanced and stage IV as advanced or metastatic breast cancer (https://www.cancerresearchuk.org/about-cancer/breast-cancer/stages-types-grades/about-breast-cancer-staging-grades).

#### Overall findings

For each dataset, we first split the gene and protein matrices into sub-matrices based on pathway information from the Kyoto Encyclopedia of Genes and Genomes (KEGG) database using the ‘KEGGREST’ package in R. The genes and proteins in our datasets belonged to 268 KEGG pathways. The number of identified genes and proteins was 246 and 83 in the TCGA while 451 and 117 in the CPTAC dataset. Although the protein assays were done on different platforms for the two datasets (reverse phase protein array or RPPA in TCGA and mass spectrometry or MS in CPTAC), interestingly, we could validate 106 genes and 65 proteins in both datasets. Further, analyses of these genes and proteins in the significant pathways revealed 64 combinations with indirect mediation effect of the genes through proteins on the disease status with no direct effect of the genes found on the same.

### Validation of the findings from three disease-gene association databases

We searched genes associated with breast cancer disease on DisGeNET v7.0^30^, KEGG ^31^, and eDGAR ^32^. Since the disease types in the two studies included invasive carcinoma, we searched for “invasive ductal carcinoma” and “invasive lobular carcinoma” disease types on DisGeNET (as it includes several sub-class for breast cancer) while “breast cancer” for the other databases. On eDGAR, 4 ioSearch identified genes (BRCA2, ESR1, BRIP1, CDH1) could be validated. Genes BRIP1 and CDH1 could not be identified in the only gene-based analysis. KEGG database validated 6 genes namely, KIT, PGR, BRIP1, BRCA2, CCND1, and EGFR out of which gene-based tests missed the first three. Out of the 591 unique genes identified in one or both of the datasets, 495 could be validated from DisGeNET when we searched for disease types containing terms (1) “breast” and “cancer” or “carcinoma” or ‘papilloma’, (2) “carcinogenesis” or “tumor initiation” or “tumor progression” or “tumor promotion”, (3) “neoplasm” and “malignant” or “mammary” or “metastasis” or “breast”, and (4) neoplasms. 154 out of 495 (Table 5 and Table *S*1) DisGeNET validated genes were missed by only gene-based tests. To find if the identified genes have been implicated via experimental approaches, we searched the literature and found a list of interesting results. We also found some crucial observations from among the genes that could not be validated by any database. Some of the results given in Table 5 with gene/protein expression in the bold case were only identified by ioSearch but missed by single-omics analyses. Because cancer is a complex disease, as expected the number of identified genes is quite high. Moreover, we aimed at selecting the top 5 genes and 3 proteins from each pathway to explore further functional relations in our model. Reducing or increasing this number is expected to vary the number of biomarkers. Conservative results are possible if the number of top omics to be selected is reduced.

**Table 5.** List of significant gene and protein expressions identified by ioSearch belonging to some important pathways. Column named ‘Both’ indicates omics identified in both TCGA and CPTAC datasets, and Columns named ‘TCGA’ and ‘CPTAC’ indicate biomarkers exclusively identified in that dataset. Protein/gene expressions in bold indicate these omics are only identified by ioSearch but missed by single omics analyses

| Pathways | Omic Type | Both | TCGA | CPTAC |
| --- | --- | --- | --- | --- |
| AMPK signaling pathway<br>(path:hsa04152) | Genes | **PRKAA2**<sup>33</sup> | **IGF1R**<sup>34</sup>, **IRS1**<sup>35</sup>,<br>CCND1, FBP1 | PPARG,<br>PPARGC1A,<br>PPP2R2B,<br>PPP2R3B |
|  | Proteins | AKT1S1 | PIK3CA, TSC1 | FOXO3, RPS6KB1 |
| Tight junction signaling pathway<br>(path:hsa04530) | Genes | PPP2R2B | **CD1C**<sup>53</sup>,<br>**CLDN1**<sup>54</sup>,<br>CLDN11, CLDN8 | BVES, CGN,<br>MICALL2,<br>**PRKAA2**<sup>33</sup> |
|  | Proteins |  | CCND1,<br>**ERBB2**<sup>36</sup>, **SRC** | MAPK9, PCNA,<br>VASP |
| Transcriptional misregulation in cancer<br>(path:hsa05202) | Genes | **NGFR**<sup>39</sup> | **BCL11B**<sup>41</sup>,<br>BCL2A1, IL6,<br>**PROM1**<sup>40</sup> | GADD45G, MMP9,<br>PPARG, RUNX1T1 |
|  | Proteins | TP53 | ATM, IGFBP3 | MITF, PTK2 |
| Antigen processing and presentation<br>(path:hsa04612) | Genes | KLRC4, KLRD1,<br>**TAP1**<sup>43</sup> | CTSS, IFI30 | RFXAP, **TAP2**<sup>43</sup> |
|  | Proteins | HSPA5 | CREB1, HSPA1A | **CANX**,<br>HLA-DQA1 |
| Nucleotide metabolism<br>(path:hsa01232) | Genes | RRM2 | **AK7**, CTPS1,<br>ENPP1, UPP1 | CDA, GMPR,<br>LACC1, TK2 |
|  | Proteins | **RRM1** |  |  |
| Biosynthesis of nucleotide sugars<br>(path:hsa01250) | Genes | HK2 | GNPNAT1,<br>**PMM2**, **UGDH**,<br>**GFPT1** | FPGT, GALE,<br>GALK2, GMPPA |
|  | Proteins | **HK1**, **HK2**<sup>45</sup>,<br>PGM1 |  |  |
| Dilated cardiomyopathy<br>(path:hsa05414) | Genes | PLN | **ADCY2**, **IGF1**,<br>LAMA2, TGFB2 | ADCY4, ITGA8,<br>**CACNA2D3**,<br>**CACNB4** |
|  | Proteins | **ACTB**, ITGA2,<br>ITGB1 |  |  |
| Type I diabetes mellitus<br>(path:hsa04940) | Genes | CD28, CD86,<br>**FASLG**, **IL1B**<sup>47</sup>,<br>PTPRN2 |  |  |
|  | Proteins | HLA-DQA1,<br>HSPD1 |  |  |

### Interesting observations

Most of the biomarkers identified using ioSearch have already been experimentally validated in different studies related to breast cancer metastasis, tumor progression, tumorigenesis etc. Some of the identified biomarkers occurring in important pathways are displayed in Table 5 and the rest in Table *S*1. Few of the biomarkers in Table 5 have also been held responsible in experimental studies for triggering interactions with other biomarkers within the pathway due to oncogenic signaling. This provides a validation that ioSearch is able to identify interacting biomarkers associated with oncogenic signaling. In the AMPK signaling pathway (in Table 5), we identified PRKAA2 gene expression and AKT1S1(PRAS40) protein expression in both TCGA and CPTAC to be associated with the phenotype of interest. As shown in Table 5, in the same pathway we also identified IRS1, IGF1R, CCND1, TSC1 etc. genes. But some of the important genes such as PRKAA2, IRS1 and IGF1R were unidentified by single-omics analyses. Each of the extra genes that ioSearch identified has been implicated by various experimental studies to have a prominent role in cancer. Evidence suggests that PRKAA2 play important role in the alteration of CCND1 expression as well as AMPK growth control and apoptotic signaling ^33^, in head and neck cancer PRAS40 phosphorylation plays an important role in IGF1R-based therapeutic resistance to EGFR inhibition ^34^, and the effect of tumor suppressor gene TSC1 in controlling insulin-PI3K signaling via regulation of IRS proteins ^35^. So, this method not only finds interacting multi-omics biomarkers but also captures weaker signals that are missed by traditional single-omics analyses. Again in the tight junction pathway (in Table 5), we identified expressions of CD1C, CLDN1, ERBB2, SRC and PRKAA2 but were missed by the single-omics tests. Of these, ERBB2/HER2 is known to be one of the most important metastasis-related biomarkers for breast cancer ^36^. Moreover, ERBB2 is related to the activation and further functioning of SRC. In addition, overexpression of SRC increases the response of EGFR-mediated processes. Thus, ERBB2 interacts with SRC to play a key role in the tumor progression of certain late-stage breast cancers ^37^. Other studies suggest suppression of CLDN1, a major component of tight junction significantly inhibits tumor growth ^38^. In pathway transcriptional misregulation in cancer, we exclusively identified NGFR ^39^, PROM1^40^ and BCL11B ^41^ genes related to tumor growth or metastasis, along with evidence supporting co-expression of the first two in some cancers ^42^. In both datasets ioSearch identified genes such as TAP1/2^43^, RRM1^44^, HK1/2^45^, ACTB ^46^, IL1B ^47^, SH2D1A ^48^ etc. that are directly related to breast cancer stages or metastasis. Other genes such as UGDH ^49^, IGF1^50^ etc. were identified in either one of the datasets belonging to the pathways of the above genes and are also related to breast cancer metastasis and/or disease progression. Further, the literature survey suggests some genes such as PABPC3^51^, PKIA ^52^ etc. (in Table *S*1) etc. that are unidentified by any disease-gene association databases and single-omics analyses are strongly related to tumorigenesis.

## Discussion

Biomarkers are important predictors of disease onset/progression and hence play a vital role in the prediction of patient survival and/or response to therapy. However, the key challenge in finding biomarkers for complex diseases is, decoding the intricate interplay of multiple omics data. But, combining results from single-omics analyses of multiple assays tends to overlook signals from omics with moderate to weak effect sizes and remains limited to the identification of strongest signals due to multiple testing burdens. On the other hand, multi-omics data analysis has the advantage of accumulating complementary information that might identify these missed out signals. Although information from multiple omics data intuitively provides substantial information on the disease compared to a single source of data, it is difficult to analyse such correlated information from multiple omics together due to the differences in the data structure emerging from different assays, less sample size than the number of features under study etc. In such circumstances, feature selection methods for dimension reduction are often employed. But almost all the methods focus on subtype classification either via supervised or unsupervised learning methods. Biomarker identification is looked upon as a by-product in some of these methods. The reason is that it is difficult to biologically interpret the meaning of significant features in terms of the original variables. The regression framework although provides better interpretation fails when the number of variables greatly outnumbers the sample size and/or multicollinearity arises. Even the recent technological developments that paved the way for the generation of individual-specific multi-omics data are not enough. Thus, it is essential to develop efficient algorithms that are able to tackle such situations. It is important to note that, biological knowledge such as pathway information or other functional annotation might provide crucial insights. This information may be used to meaningfully reduce dimensions which in turn might aid in the interpretation of features.

Keeping all these in mind, we proposed ioSearch for the identification of biomarkers that may differentiate between two disease statuses by integrating two omics assays with continuous measurements. To demonstrate our method we selected two omics viz. transcriptomics and proteomics from two publicly available breast cancer datasets. The reason behind selecting these omics is driven by the fact that protein synthesized from mRNAs is oftentimes a primary downstream output of oncogenic signaling. Our method is able to explore the inter-relation between omics that are functionally related (eg. belonging to the same pathway) and also the overall effect of each omics type on the disease. In doing so, ioSearch reduces multiple testing burdens thus allowing the capture of weaker signals with smaller effect sizes, which might otherwise be lost if analysed individually. The inclusion of pathway information from the KEGG database facilitates meaningful dimension reduction and also in the exploration of long-range functional relations. This prior information in addition to our comprehensive testing algorithm allows the identification of significant multi-omics markers, by analysing the entire multi-omics datasets as opposed to gene-wise analysis. To handle multicollinearity in each pathway and for better interpretation of the features, we have used the PCR framework and SPCA. Moreover, ioSearch is a time-cost efficient algorithm that provides easily interpretable p-values.

Simulation results confirm the consistency of the method, controlled type I error and high statistical power compared to existing methods. We observe that the statistical power is independent of the number of pathways or functional annotations considered but is dependent on the percentage of the true significant sets. Although the latter is expected for an efficient algorithm, the former result shows that our method will work similarly even when prior information varies. Starting with simple scenarios we moved on to more complex scenarios of interactions to demonstrate how our method will work in real datasets. To depict some complexities that might be common in real datasets we simulated between-omics interactions with varying combinations of magnitude and directions. ioSearch produced satisfactory results under every simulation setup.

On application of ioSearch to the protein and gene expression data of nearly 800 and 100 breast cancer patients from TCGA and CPTAC databases, we identified a list of genes and proteins most of which are reported to have interacting functional implications across independent experimental studies. Some of them were identified in both datasets but missed by commonly used single omics analyses. Most of the identified biomarkers are already published and interestingly they are shown to be associated with our phenotype of interest. Moreover, we found evidence from multiple experimental approaches that confirm biological interactions among the identified biomarkers pertaining to the same pathway. Thus, ioSearch is able to identify interacting biomarkers associated with the phenotype of interest. It can be used as a tool for either biomarker identification by wet-lab researchers on the generated data or to narrow down the list of aberrantly behaving omics from public data repositories that are associated with the phenotype of interest, for further functional experiments.

