## Supplementary Table 1 for "ioSearch: a tool for searching disease-associated interacting omics; application on breast cancer data"

Sarmistha Das and Deo Kumar Srivastava
Department of Biostatistics, St. Jude Children's Research Hospital, Memphis, TN

S 1: List of significant gene and protein expressions identified by ioSearch belonging to all the significant pathways.

| Pathways | Omic Type | Both | TCGA | CPTAC |
|---|---|---|---|---|
| AMPK signaling pathway (path:hsa04152) | Genes | **PRKAA2**[1] | **IGF1R**[2], **IRS1**[3], CCND1, FBP1 | PPARG, PPARGC1A, PPP2R2B, PPP2R3B |
| | Proteins | AKT1S1 | PIK3CA, TSC1 | FOXO3, RPS6KB1 |
| Tight junction signaling pathway (path:hsa04530) | Genes | PPP2R2B | **CD1C**[4], **CLDN1**[5], CLDN11, CLDN8 | BVES, CGN, MICALL2, **PRKAA2**[1] |
| | Proteins | | CCND1, **ERBB2**[6], **SRC** | MAPK9, PCNA, VASP |
| Transcriptional misregulation in cancer (path:hsa05202) | Genes | **NGFR**[7] | **BCL11B**[8], BCL2A1, IL6, **PROM1**[9] | GADD45G, MMP9, PPARG, RUNX1T1 |
| | Proteins | TP53 | ATM, IGFBP3 | MITF, PTK2 |
| Antigen processing and presentation (path:hsa04612) | Genes | KLRC4, KLRD1, **TAP1**[10] | CTSS, IFI30 | RFXAP, **TAP2**[10] |
| | Proteins | HSPA5 | CREB1, HSPA1A | **CANX**, HLA-DQA1 |
| Nucleotide metabolism (path:hsa01232) | Genes | RRM2 | **AK7**, CTPS1, ENPP1, UPP1 | CDA, GMPR, LACC1, TK2 |
| | Proteins | **RRM1** | | |
| Biosynthesis of nucleotide sugars (path:hsa01250) | Genes | HK2 | GNPNAT1, **PMM2**, **UGDH**, **GFPT1** | FPGT, GALE, GALK2, GMPPA |
| | Proteins | **HK1**, **HK2**[11], PGM1 | | |

| | | | | |
|---|---|---|---|---|
| Dilated cardiomyopathy (path:hsa05414) | Genes | PLN | **ADCY2**, **IGF1**, LAMA2, TGFB2 | ADCY4, ITGA8, **CACNA2D3**, **CACNB4** |
| | Proteins | **ACTB**, ITGA2, ITGB1 | | |
| Type I diabetes mellitus (path:hsa04940) | Genes | CD28, CD86, **FASLG**, **IL1B**[12], PTPRN2 | | |
| | Proteins | HLA-DQA1, HSPD1 | | |
| Tyrosine metabolism (path:hsa00350) | Genes | **AOX1**, **HGD**, **MAOA** | ALDH3B2, **MAOB** | **AOC2**, AOC3 |
| | Proteins | MIF | | |
| Calcium signaling pathway (path:hsa04020) | Genes | **CD38**, EGFR, **FGF7**, **PDGFRA**, PRKCB | CAMK4, HGF, MCOLN2, RYR1, TACR2 | |
| | Proteins | **ERBB2** | **PDGFRB**, VDAC1 | FGF2, KDR |
| Alcoholic liver disease (path:hsa04936) | Genes | **FASLG**, IL6, PPARGC1A | C3, CXCL2 | IL17RA, **PRKAA2** |
| | Proteins | | **GSK3B**, **MAPK14**, RIPK1 | CTNNB1, FOXO3, IRF3 |
| Neomycin (path:hsa00524) | Genes | HK1, HK2 | | |
| | Proteins | **HK1**, **HK2** | | |
| Ubiquinone and other terpenoid-quinone biosynthesis (path:hsa00130) | Genes | **COQ3**, COQ5, **COQ7**, NQO1 | **VKORC1L1** | COQ2 |
| | Proteins | NQO1 | | |
| Purine metabolism (path:hsa00230) | Genes | PDE11A | GUCY1A2, **ADCY1**, **AK7**, ENPP1 | **PDE1C**, PDE3B, **PDE7B**, ENPP4 |
| | Proteins | PGM1, **RRM1** | | |
| Cysteine and methionine metabolism (path:hsa00270) | Genes | **CBS**, PHGDH, **PSAT1** | **LDHB**, DNMT3B | **CDO1**, LACC1 |
| | Proteins | GCLC, GCLM, PHGDH | | |
| Primary immunodeficiency (path:hsa05340) | Genes | **CD3D**, CD3E, **CD79A**, IL7R | LCK | **JAK3** |

| | | | | |
|---|---|---|---|---|
| | Proteins | **CD4**, ZAP70 | | |
| Pyrimidine metabolism (path:hsa00240) | Genes | ENPP1, RRM2 | CMPK2, CTPS1, UPP1 | CDA, NT5E, TK2 |
| | Proteins | **RRM1** | | |
| Viral life cycle - HIV-1 (path:hsa03250) | Genes | APOBEC3D, **APOBEC3G**, CCR5 | APOBEC3A, **MX1** | **APOBEC3F**, MAP1A |
| | Proteins | **CD4** | | |
| Base excision repair (path:hsa03410) | Genes | POLD1, POLE | LIG1, PCNA, **UNG** | **MPG**, PARP3, POLL |
| | Proteins | **PARP1**, PCNA, XRCC1 | | |
| Cytokine-cytokine receptor interaction (path:hsa04060) | Genes | | **CCL13**, CCL19, CXCL13, **CXCL9**, IL7R | ACVR1C, **CXCR3**, **FASLG**, IL10, IL18RAP |
| | Proteins | **CD4** | | AKT2, MAPK9, PAK1 |
| Notch signaling pathway (path:hsa04330) | Genes | LFNG, MAML2, TLE6 | HEY2, **TLE4** | DVL1, MAML3 |
| | Proteins | HES1, NOTCH1, **NOTCH3** | | |
| Hematopoietic cell lineage (path:hsa04640) | Genes | **CD38**, CD3G, **MS4A1** | CD3E, IL7R | **CD1C**, **CR1** |
| | Proteins | **CD38** | CD44, KIT | ITGA2, TFRC |
| Natural killer cell mediated cytotoxicity (path:hsa04650) | Genes | CD244, **FASLG**, PRKCB, **SH2D1A** | LCK | KLRD1 |
| | Proteins | | BID, MAPK1, PIK3CA | ARAF, NRAS, PAK1 |
| Th17 cell differentiation (path:hsa04659) | Genes | CD3G, **IRF4**, TBX21 | **CD3D**, CD3E | **EBI3**, IL6 |
| | Proteins | | MAPK1, **MAPK14**, RELA | MAPK9, SMAD3, ZAP70 |
| Fc gamma R-mediated phagocytosis (path:hsa04666) | Genes | **PIP5K1B**, PRKCA, PRKCB | **PLA2G4A**, **PTPRC** | PRKCE, WASF3 |
| | Proteins | PRKCD | MAPK1, PIK3CA | PAK1, RPS6KB1 |

| | | | | |
|---|---|---|---|---|
| Circadian entrainment (path:hsa04713) | Genes | **ADCYAP1R1**, GUCY1A2 | **ADCY2**, **FOS**, PRKCB | ADCY4, PRKG2, RYR1 |
| | Proteins | CREB1, MAPK1, PRKCA | | |
| Thermogenesis (path:hsa04714) | Genes | **PRKAA2**, RPS6KA6 | **ADCY1**, PLIN1, ZNF516 | COX4I2, PPARG, PPARGC1A |
| | Proteins | | ATP5PD, **RPS6**, TSC1 | AKT1S1, NRAS, RPS6KB1 |
| Adipocytokine signaling pathway (path:hsa04920) | Genes | PPARGC1A, **PRKAA2** | **ACSL4**, **IRS2**, LEPR | ACSL6, **MAPK10**, PPARA |
| | Proteins | ACSL1, IRS2 | **PTPN11** | AKT2 |
| Pancreatic secretion (path:hsa04972) | Genes | PRKCA, PRKCB | **ADCY2**, ADCY7, **CD38** | ATP1A3, RAB27B, TRPC1 |
| | Proteins | **CD38**, PRKCA | | |
| Alcoholism (path:hsa05034) | Genes | **PKIA** | FOSB, **MAOA**, **MAOB**, NTRK2 | CAMK4, CREB5, GNAO1, SHC2 |
| | Proteins | NRAS | **H3C1**, MAPK1 | ARAF, H2AX |
| Legionellosis (path:hsa05134) | Genes | **CR1**, CXCL2, CXCL8, IL6 | C3 | IL18 |
| | Proteins | CASP7, HSPD1 | RELA | CASP8 |
| Leishmaniasis (path:hsa05140) | Genes | **CR1**, PRKCB | C3, CYBB, **IL1B** | IL10, MAPK11, PTGS2 |
| | Proteins | MAPK1 | **MAPK14**, RELA | HLA-DQA1, ITGB1 |
| Chagas disease (path:hsa05142) | Genes | CD3G, **FASLG**, IL6 | **CD3D**, CD3E | IL10, PPP2R2B |
| | Proteins | PIK3CA | MAPK1, **MAPK14** | AKT2, MAPK9 |
| Hepatitis B (path:hsa05161) | Genes | **FASLG**, IL6, PRKCB | **EGR2**, **EGR3** | PRKCA, TGFB2 |
| | Proteins | | MAPK1, PIK3CA, **YWHAB** | ARAF, IRF3, NRAS |
| Viral myocarditis (path:hsa05416) | Genes | CD28, ICAM1, **ITGAL** | **CD40**, **RAC2** | DMD, FYN |
| | Proteins | | BID, CASP8, EIF4G1 | CAV1, CCND1, HLA-DQA1 |

| | | | | |
|---|---|---|---|---|
| Autoimmune thyroid disease (path:hsa05320) | Genes | CD28, **CTLA4**, **FASLG**, IL10 | CD86 | TG |
| | Proteins | HLA-DQA1 | | |
| Toxoplasmosis (path:hsa05145) | Genes | CCR5 | BIRC3, **IL10RA**, LAMA2, TGFB2 | GNAO1, IL10, LAMA3, PIK3R6 |
| | Proteins | | MAPK1, **MAPK14**, RELA | AKT2, HLA-DQA1, MAPK9 |
| Yersinia infection (path:hsa05135) | Genes | IL6, NLRP3 | **IL1B**, LCK, **PIP5K1B** | IL10, RPS6KA6, WIPF3 |
| | Proteins | | MAPK1, **MAPK14**, PIK3CA | AKT2, IRF3, ZAP70 |
| Parathyroid hormone synthesis (path:hsa04928) | Genes | PRKCA, PRKCB | **ADCY2**, EGFR, PDE4B | **ADCY1**, PLCB4, PTH1R |
| | Proteins | MAPK1 | BRAF, CREB1 | ARAF, RAF1 |
| Non-alcoholic fatty liver disease (path:hsa04932) | Genes | IL6 | **COX6C**, CXCL8, **FASLG**, **FOS** | COX4I2, PPARG, **PRKAA2**, SREBF1 |
| | Proteins | | BID, CASP7, PIK3CA | AKT2, INSR, MAPK9 |
| Age-rage signaling pathway in diabetic complications (path:hsa04933) | Genes | IL6, PRKCB | **COL3A1**, SELE, TGFB2 | COL4A5, PLCB4, PRKCA |
| | Proteins | PRKCD | MAPK1, PIK3CA | AKT2, NRAS |
| Growth hormone synthesis (path:hsa04935) | Genes | PRKCB | **ADCY2**, **FOS**, **IGF1**, **SOCS2** | **ADCY1**, PLCB4, PRKCA, SHC2 |
| | Proteins | NRAS | MAPK1, PIK3CA | AKT2, MAPK9 |
| Vasopressin-regulated water reabsorption (path:hsa04962) | Genes | DYNC1I1, PRKACB | **AQP3**, ARHGDIB, **CREB3L1** | ADCY9, CREB5, DYNC2H1 |
| | Proteins | CREB1 | | |
| Fatty acid biosynthesis (path:hsa00061) | Genes | ACACB, ACSL1, ACSL6, FASN | **ACSL4** | ACSF3 |
| | Proteins | ACSL1, FASN | | |
| Fatty acid degradation (path:hsa00071) | Genes | ACSL6 | **ACADSB**, ACSL1, **ACSL4**, ECI2 | ACADS, EHHADH, ALDH7A1, CPT1A |

| | | | | |
|---|---|---|---|---|
| | Proteins | ACSL1 | | |
| Oxidative phosphorylation (path:hsa00190) | Genes | ATP6V0D2 | **COX6C**, **NDUFA11**, NDUFS6, NDUFS8 | ATP6V0E2, ATP6V1B1, **COX4I2**, NDUFS7 |
| | Proteins | ATP5PD, COX4I1, SDHA | | |
| NF-kappa B signaling pathway (path:hsa04064) | Genes | PRKCB | BCL2A1, **CCL13**, CCL19, LCK | CARD10, CXCL2, GADD45G, PTGS2 |
| | Proteins | TRIM25 | ATM, RIPK1 | SYK, ZAP70 |
| FoxO signaling pathway (path:hsa04068) | Genes | IL6 | EGFR, **IGF1**, IL7R, TGFB2 | CCNB3, **FASLG**, FOXO6, IL10 |
| | Proteins | | ATM, MAPK1, PTEN | ARAF, FOXO3, NRAS |
| Mitophagy (path:hsa04137) | Genes | | **ATG9A**, FIS1, HIF1A, **MAP1LC3A**, RHOT2 | FOXO3, **MAPK10**, **MITF**, PGAM5, RRAS2 |
| | Proteins | NRAS | AMBRA1, **SRC** | FOXO3, MITF |
| Phagosome (path:hsa04145) | Genes | | C3, **CD209**, CYBB, MRC1, **THBS2** | ATP6V0D2, DYNC1I1, OLR1, TLR6, TUBA8 |
| | Proteins | TFRC | **CANX**, HLA-DQA1 | ITGA2, ITGB1 |
| Adrenergic signaling in cardiomyocytes (path:hsa04261) | Genes | ADRB2, PPP2R2B | **ADCY2**, **CACNB4**, PLN | ATP1A3, PIK3R6, PRKCA |
| | Proteins | CREB1, MAPK1 | **MAPK14** | AKT2 |
| Vascular smooth muscle contraction (path:hsa04270) | Genes | EDN2 | **ACTG2**, **ADCY2**, **EDN1**, **PLA2G4A** | AVPR1A, PLCB4, PRKCA, PRKCB |
| | Proteins | MYH9, PRKCD | MAPK1 | ARAF |
| Hippo signaling pathway (path:hsa04390) | Genes | BMP2 | BMPR1B, CCND1, PARD6B, **WNT11** | AXIN2, FZD10, GLI2, PPP2R2B |
| | Proteins | | **GSK3B**, WWTR1, **YWHAB** | CCND1, CTNNB1, SMAD3 |
| Cell adhesion molecules (path:hsa04514) | Genes | SELP | CD2, **CTLA4**, **PTPRC**, **SELL** | CNTNAP2, ITGA8, NRXN2, NTNG2 |
| | Proteins | CLDN7 | **CD4**, HLA-DQA1 | ITGB1, PECAM1 |

| | | | | |
|---|---|---|---|---|
| C-type lectin receptor signaling pathway (path:hsa04625) | Genes<br>Proteins | CCL22, IL6, PTGS2<br>PRKCD | **CD209**, **CLEC4E**<br>MAPK1, PIK3CA | IL10, NLRP3<br>NRAS, PAK1 |
| Fc epsilon RI signaling pathway (path:hsa04664) | Genes<br><br>Proteins | <br><br>NRAS | BTK, **FCER1A**, INPP5D, **LCP2**, **RAC2**<br><br>MAPK1, PIK3CA | LAT, **MAPK10**, MAPK11, PRKCA, RAC3<br><br>AKT2, SYK |
| Th1 and Th2 cell differentiation (path:hsa04658) | Genes<br><br>Proteins | CD3G, TBX21 | **CD3D**, CD3E, LCK<br><br>MAPK1, **MAPK14**, RELA | IL12RB1, **JAK3**, MAML3<br><br>MAPK9, NOTCH1, ZAP70 |
| TNF signaling pathway (path:hsa04668) | Genes<br><br>Proteins | CXCL2, PTGS2<br><br>RIPK1 | **CCL5**, IL6, SELE<br><br>MAPK1, PIK3CA | LIF, MMP9, RIPK3<br><br>AKT2, MAPK9 |
| Intestinal immune network for IgA production (path:hsa04672) | Genes<br><br>Proteins | CD28, IL10, IL6<br><br>HLA-DQA1 | CD86, **TNFSF13B** | ICOSLG, IL15 |
| Glutamatergic synapse (path:hsa04724) | Genes<br><br>Proteins | PRKCB<br><br>GLS, MAPK1, PRKCA | **GNG4**, **PLA2G4A**, PLCB4, PRKCA | GNAO1, GRM8, SHANK2, TRPC1 |
| Cholinergic synapse (path:hsa04725) | Genes<br><br>Proteins | CAMK4, PRKCB<br><br>NRAS, PIK3CA | **ADCY2**, KCNQ5, PRKCA<br><br>MAPK1 | CHRNA7, KCNJ12, KCNJ18<br><br>AKT2 |
| Serotonergic synapse (path:hsa04726) | Genes<br><br>Proteins | <br><br>MAPK1, NRAS | **ALOX15B**, **CYP4X1**, **GNG4**, **MAOA**, **MAOB**<br><br>BRAF | GNAO1, KCND2, PRKCB, PTGS2, TRPC1<br><br>ARAF |
| Melanogenesis (path:hsa04916) | Genes<br><br>Proteins | PRKCB<br><br>NRAS | **ADCY2**, **EDN1**, **KIT**, PRKCA<br><br>KIT, MAPK1 | FZD10, FZD2, GNAO1, WNT2<br><br>CTNNB1, MITF |
| Salmonella infection (path:hsa05132) | Genes<br><br>Proteins | FLNC, IL6 | BIRC3, **IL1B**, **PTPRC**<br><br>MAPK1, PIK3CA, RIPK1 | DYNC1I1, NLRP3, WASF3<br><br>AKT2, CTNNB1, PAK1 |

| | | | | |
|---|---|---|---|---|
| Pertussis (path:hsa05133) | Genes<br>Proteins | IL6<br>CASP7 | **C1S**, C3, **IL1B**, **IRF8**<br>MAPK1, **MAPK14** | IL10, MAPK11, NLRP3, TICAM1<br>IRF3, MAPK9 |
| Graft-versus-host disease (path:hsa05332) | Genes<br>Proteins | <br>HLA-DQA1 | CD28, **FASLG**, **IL1B**, IL6, KLRD1 | |
| Arrhythmogenic right ventricular cardiomyopathy (path:hsa05412) | Genes<br>Proteins | LAMA2<br>GJA1 | CDH2, **GJA1**, **ITGA11**, ITGA2<br>**ACTB**, ITGB1 | **CACNA2D3**, **CACNB4**, ITGA8, **ITGB3**<br>CTNNB1, ITGA2 |
| Lipid and atherosclerosis (path:hsa05417) | Genes<br>Proteins | IL6 | **CCL5**, CXCL2, **FASLG**, SELE<br>HSPD1, MAPK1, PIK3CA | NLRP3, POU2F2, PPARG, PRKCA<br>AKT2, IRF3, NRAS |
| Acute myeloid leukemia (path:hsa05221) | Genes<br>Proteins | <br>NRAS | BCL2A1, CCND1, **CSF1R**, **KIT**, **PIM1**<br>MAPK1, PIK3CA | ITGAM, PER2, RPS6KB2, RUNX1T1, SPI1<br>ARAF, RPS6KB1 |
| Inflammatory bowel disease (path:hsa05321) | Genes<br>Proteins | IL18RAP, IL6, TBX21<br>HLA-DQA1, RELA, SMAD3 | **IL1B**, STAT4 | IL10, **IL18R1** |
| Rheumatoid arthritis (path:hsa05323) | Genes<br>Proteins | IL6<br>HLA-DQA1 | **CCL5**, **CTLA4**, CXCL2, **IL1B** | ANGPT1, ATP6V0D2, , ATP6V1B1, CD28 |
| Pathogenic Escherichia coli infection (path:hsa05130) | Genes<br>Proteins | IL6, NLRP3<br>RIPK1 | CLDN1, FASLG, NCKAP1L<br>MAPK1, PTPN11 | MYO5C, WASF3, WIPF3<br>MAPK9, PAK1 |
| Allograft rejection (path:hsa05330) | Genes<br>Proteins | CD28, **CD40**, CD86, **FASLG**, IL10<br>HLA-DQA1 | | |

| | | | | |
|---|---|---|---|---|
| D-Amino acid metabolism (path:hsa00470) | Genes | DDO | | |
| | Proteins | GLS | | |
| Cellular senescence (path:hsa04218) | Genes | | **CCNE1**, **CDC25A**, CXCL8, **ITPR1**, MYBL2 | CCNB2, CCNB3, GADD45G, IL6, TRPV4 |
| | Proteins | | MAPK1, PTEN, TSC1 | CHEK2, FOXO3, NRAS |
| Relaxin signaling pathway (path:hsa04926) | Genes | | **ADCY2**, **COL3A1**, EGFR, **FOS**, **MMP2** | COL4A5, EDNRB, GNAO1, MMP9, SHC2 |
| | Proteins | | MAPK1, **MAPK14**, PIK3CA | AKT2, MAPK9, NRAS |
| Platelet activation (path:hsa04611) | Genes | | **ADCY2**, ADCY7, **COL3A1**, PIK3CG, **PLA2G4A** | ADCY4, GUCY1A2, P2RY1, PLCB3, PRKG2 |
| | Proteins | | MAPK1, **MAPK14**, PIK3CA | AKT2, ITGA2, SYK |
| Amino sugar and nucleotide sugar metabolism (path:hsa00520) | Genes | | **CHIT1**, **GFPT2**, **NPL**, **RENBP**, **UGDH** | |
| | Proteins | | **HK1**, **HK2**, PGM1 | |
| Alanine (path:hsa00250) | Genes | | ASRGL1, ASS1, **FOLH1**, **GFPT2**, **GLUL** | |
| | Proteins | | ASNS, GLS, GLUD1 | |
| Glycine (path:hsa00260) | Genes | | **CBS**, GAMT, **MAOA**, PHGDH, **PSAT1** | |
| | Proteins | | PHGDH | |
| Pyruvate metabolism (path:hsa00620) | Genes | | | ACACB, ALDH3A2, ALDH7A1, **ME1**, ME2 |
| | Proteins | | | ACSS2, DLAT, PDHA1 |
| Pyruvate metabolism (path:hsa00620) | Genes | | **AOX1**, **CD38**, NNMT, NT5E, **NUDT12** | |

| | | | | |
|---|---|---|---|---|
| | Proteins | | **CD38** | |
| Nitrogen metabolism (path:hsa00910) | Genes<br>Proteins | | **CA12**, CA2, CPS1, GLUD2, **GLUL**<br>GLUD1 | |
| Biosynthesis of amino acids (path:hsa01230) | Genes<br>Proteins | | ASS1, **CBS**, **PFKP**, PHGDH, **PSAT1**<br>ASNS, **ENO1**, PHGDH | |
| Antifolate resistance (path:hsa01523) | Genes<br>Proteins | | | ABCC4, **ABCG2**, FOLR2, IL6, MTHFR<br>RELA |
| RNA degradation (path:hsa03018) | Genes<br>Proteins | | BTG2, BTG3, **PABPC3**, PABPC5, **PFKP**<br>**ENO1**, HSPA9, HSPD1 | |
| PPAR signaling pathway (path:hsa03320) | Genes<br>Proteins | | | ACOX2, ACSL6, PLIN5, PPARA, PPARG<br>ACSL1, **PDPK1** |
| cAMP signaling pathway (path:hsa04024) | Genes<br>Proteins | | | **ADCYAP1R1**, ATP1A3, CAMK4, EDN2, PTCH1<br>AKT2, MAPK9, PAK1 |
| Cell cycle (path:hsa04110) | Genes<br>Proteins | | **CCNE1**, CDC20, **CDC25A**, PLK1, TTK<br>ATM, ATR, **YWHAB** | |
| Peroxisome (path:hsa04146) | Genes<br>Proteins | | ACOX2, ACSL6, **CRAT**, **CROT**, MPV17L<br>ACSL1, **PRDX1**, **SOD2** | |
| Osteoclast differentiation (path:hsa04380) | Genes | | **CSF1R**, **IL1B**, LCK, **LILRB1**, SIRPB1 | |

| | | | | |
|---|---|---|---|---|
| | Proteins | | MAPK1, PIK3CA, **MAPK14** | |
| RIG-I-like receptor signaling pathway (path:hsa04622) | Genes | | CXCL10, CXCL8, **IFIH1**, **ISG15**, **MAPK10** | |
| | Proteins | | **ATG5**, RIPK1, TRIM25 | |
| B cell receptor signaling pathway (path:hsa04662) | Genes | | CARD11, **CD79A**, **CD79B**, **LILRB1**, PRKCB | |
| | Proteins | | MAPK1, NRAS, PIK3CA | |
| Neurotrophin signaling pathway (path:hsa04722) | Genes | | | CAMK4, **FASLG**, RPS6KA6, SH2B2, SHC2 |
| | Proteins | | | AKT2, FOXO3, NRAS |
| Insulin signaling pathway (path:hsa04910) | Genes | | | PPARGC1A, **PRKAA2**, **PYGM**, SH2B2, SHC2 |
| | Proteins | | | ARAF, NRAS, RPS6KB1 |
| Regulation of lipolysis in adipocytes (path:hsa04923) | Genes | | **ADCY1**, ADRB2, **IRS1**, NPY1R, PLIN1 | |
| | Proteins | | IRS1, IRS2, PIK3CA | |
| Salivary secretion (path:hsa04970) | Genes | | **ADCY2**, **CD38**, **LYZ**, PRKCA, PRKCB | |
| | Proteins | | **CD38**, PRKCA | |
| Cholesterol metabolism (path:hsa04979) | Genes | | | CYP27A1, **LIPG**, LRP1, **LRP2**, NCEH1 |
| | Proteins | | | VDAC1 |
| Staphylococcus aureus infection (path:hsa05150) | Genes | | **C1S**, C3, KRT17, **KRT23**, SELP | |
| | Proteins | | HLA-DQA1 | |
| Gastric cancer (path:hsa05226) | Genes | | EGFR, **FGF7**, MET, TGFB2, WNT5A | |
| | Proteins | | **ERBB2**, FGF2, MAPK1 | |

| | | | | |
|---|---|---|---|---|
| Hypertrophic cardiomyopathy (path:hsa05410) | Genes | | DMD, **IGF1**, IL6, LAMA2, TGFB2 | |
| | Proteins | | **ACTB**, ITGA2, ITGB1 | |
| Pathways in cancer (path:hsa05200) | Genes | | | GLI2, PRKCB, PTCH1, PTGS2, RUNX1T1 |
| | Proteins | | | ARAF, NOTCH1, NRAS |
| MicroRNAs in cancer (path:hsa05206) | Genes | | | ERBB3, FZD3, MMP9, PRKCB, PTGS2 |
| | Proteins | | | MCL1, NOTCH1, NRAS |
| Chemical carcinogenesis - reactive oxygen species (path:hsa05208) | Genes | | | COX4I2, EPHX3, GSTM1, HGF, PRKD1 |
| | Proteins | | | ARAF, FOXO3, NRAS |
| Chronic myeloid leukemia (path:hsa05220) | Genes | | | CBL, GADD45G, SHC2, TGFB2, TGFB3 |
| | Proteins | | | AKT2, ARAF, NRAS |
| Regulation of actin cytoskeleton (path:hsa04810) | Genes | | | BCAR1, FGF1, ITGA8, LPAR1, PDGFD |
| | Proteins | | | ARAF, NRAS, PAK1 |
| Axon guidance (path:hsa04360) | Genes | | | NTNG2, PTCH1, SEMA3A, SRGAP1, TRPC4 |
| | Proteins | | | NRAS, PAK1, PTK2 |
| ErbB signaling pathway (path:hsa04012) | Genes | | | ERBB3, NRG1, PRKCA, PRKCB, SHC2 |
| | Proteins | | | ARAF, NRAS, RPS6KB1 |
| Sphingolipid signaling pathway (path:hsa04071) | Genes | | | ACER2, ADORA1, ASAH2, PPP2R2B, PRKCB |

| | | | | |
|---|---|---|---|---|
| | Proteins | | | AKT2, NRAS, TP53 |
| Metabolic pathways (path:hsa01100) | Genes | | | ATP6V0D2, DGKG, OXCT2, PDE11A, RIMKLA |
| | Proteins | | | ASNS, GCLM, NQO1 |
| Fatty acid metabolism (path:hsa01212) | Genes | | | ACSF3, ACSL6, EHHADH, ELOVL4, ELOVL7 |
| | Proteins | | | ACSL1, FASN |
| Lysine degradation (path:hsa00310) | Genes | | | AADAT, AASS, ALDH7A1, KMT2C, SETD1B |
| | Proteins | | | SETD2 |

Column named 'Both' indicates omics identified in both TCGA and CPTAC datasets, and Columns named 'TCGA' and 'CPTAC' indicate biomarkers exclusively identified in that dataset. Protein/gene expressions in bold indicate these omics are only identified by ioSearch but missed by single omics analyses
